# baredSC: Bayesian Approach to Retrieve Expression Distribution of Single-Cell

**DOI:** 10.1101/2021.05.26.445740

**Authors:** Lucille Lopez-Delisle, Jean-Baptiste Delisle

## Abstract

The number of studies using single-cell RNA sequencing (scRNA-seq) is constantly growing. This powerful technique provides a sampling of the whole transcriptome of a cell. However, the commonly used droplet-based method often produces very sparse samples. Sparsity can be a major hurdle when studying the distribution of the expression of a specific gene or the correlation between the expressions of two genes. We show that the main technical noise associated with these scRNA-seq experiments is due to the sampling (i.e. Poisson noise). We developed a new tool named baredSC, for Bayesian Approach to Retrieve Expression Distribution of Single-Cell, which infers the intrinsic expression distribution in single-cell data using a Gaussian mixture model (GMM). baredSC can be used to obtain the distribution in one dimension for individual genes and in two dimensions for pairs of genes, in particular to estimate the correlation in the two genes’ expressions. We apply baredSC to simulated scRNA-seq data and show that the algorithm is able to uncover the expression distribution used to simulate the data, even in multi-modal cases with very sparse data. We also apply baredSC to two real biological data sets. First, we use it to measure the anti-correlation between *Hoxd13* and *Hoxa11*, two genes with known genetic interaction in embryonic limb. Then, we study the expression of *Pitx1* in embryonic hindlimb, for which a trimodal distribution has been identified through flow cytometry. While other methods to analyze scRNA-seq are too sensitive to sampling noise, baredSC reveals this trimodal distribution.

## Introduction

The single-cell RNA sequencing (scRNA-seq) method allows to assess the transcriptome of individual cells [Tang et al., 2009]. Thanks to the decreasing cost and to the simplification of the protocols, this technique is more and more popular. One can distinguish two classes of methods: the plate-based methods and the droplet-based methods (also called microfluidic-based methods). The first ones often provide a better estimation of genes expression for individual cells, as the number of reads per cell is usually higher. However, the experimental procedure is more complicated and these methods typically generate data for hundreds of cells. The second ones are more widely used and provide usually sparser information (less reads per cell) but for a larger number of cells (typically 50 times more) [Svensson, 2020]. As a consequence, droplet-based methods are often preferred when complex tissues are studied. Indeed, they allow to identify more precisely pools of cells with close cellular identity (clusters) [Tanay and Regev, 2017]. Gene expression within clusters are then studied in more details which is not possible when the number of cells is too low. However, with droplet-based methods, the number of reads for a given cell is very low compared to the complexity of the transcriptome. Therefore, lowly expressed genes are not always detected, and if detected, their expression level is poorly constrained. The fact that a gene for a given cell is not detected while it is actually expressed is sometimes called ‘dropout’ [Bacher and Kendziorski, 2016; Vallejos et al., 2017; Haque et al., 2017; Svensson, 2020; Nayak and Hasija, 2021]. The origin and modelisation of the ‘dropout’ phenomenon, and more generally of the observed variance in scRNA-seq is debated, as reflected by the number of different models used in tools dedicated to differential expression analysis [Wang et al., 2019]. Here, we show that Poisson noise due to sampling explains very well the observed variance (dropout included) in droplet-based scRNA-seq. We find no evidence of more complex noise sources, such as additional dropouts, negative binomial distribution, etc.

For most common applications of scRNA-seq (clustering and identification of gene markers, projection like tSNE or UMAP) the sparsity of the data does not have a strong impact [Pollen et al., 2014]. However, the expression distribution of a given gene in a population of cells is difficult to estimate and its representation by a kernel density estimation (KDE) plot, like in the violin plot of Seurat [Butler et al., 2018] or Scanpy [Wolf et al., 2018], can be misleading. Indeed, the sampling noise strongly spread the signal, and the cells with no count appear as an artificial homogeneous sub-population, mixing cells with low and no expression.

Gene correlation has been extensively studied in bulk RNA-seq in order to build regulatory networks [Segal et al., 2003]. A recent study using scRNA-seq discovered gene covariations involving microRNA [Tarbier et al., 2020]. miRNA are small RNAs which regulate gene expression by post-transcriptional processes [Bartel, 2018]. Such results would be difficult to obtain with bulk RNA-seq, showing the power of this type of study in homogenous scRNA-seq. However, this study has been conducted with a plate-based method on a very homogeneous population (mouse embryonic stem cells). When using droplet-based data, the sparsity of the data is a major barrier to this type of analysis restricting it to genes with high expression.

Recently, Breda et al. [2021] proposed a Bayesian normalization procedure called Sanity (SAmpling-Noise-corrected Inference of Transcription activitY). This method aims at correcting the counts of each cell from the sampling noise. The procedure is fast and can be applied before clustering and projection algorithms to improve their performance. It also prevents spurious correlation between pairs of genes. However, in order to efficiently perform these corrections, Sanity uses simplifying assumptions in the modeling of the genes’ expression distributions. Here, we also propose a Bayesian approach to disentangle the intrinsic variability in gene expressions from the sampling noise. However, instead of correcting the expression level of each cell, we estimate directly the expression distribution of the population of cells. Our tool, named baredSC (Bayesian Approach to Retrieve Expression Distribution of Single-Cell), approximates the expression distribution of a gene by a Gaussian mixture model (GMM). This tool is dedicated to studies where the distribution of few genes needs to be estimated precisely. It can be used to retrieve the expression distribution of a single gene, but also to infer the joint distribution of two genes in order to study genetic interactions even when the gene expression level is low and thus the frequency of non-detection is high. Using simulated data we show that it largely outperforms the classical density/violin representation and in most cases very accurately reproduces the original distribution, both in the one-dimensional and two-dimensional cases. We also show that when multi-modal distributions are simulated it gives more accurate results than Sanity. We also use real biological data sets to illustrate the power of baredSC to assess the correlation between genes or to reveal the multi-modality of a lowly expressed gene.

## Results

### Poisson distribution is a good approximation for droplet-based single-cell RNA-seq data

We focus our analysis on droplet-based scRNA-seq as this is where the sparsity of the data is a major issue. We first evaluate the variability coming from the technique itself, including all steps of the protocol from RNA to the gene count matrix. Indeed, the source of variability in droplet-based scRNA-seq is still debated. The most obvious source of noise comes from the sampling, i.e. the fact that only a subset of the whole transcriptome is sequenced. This sampling noise is especially strong for lowly expressed genes and when the total number of reads per cell is low. In addition to the sampling noise, some specific steps of the technique, like the capture of mRNA or the amplification steps, could possibly introduce variability and/or biases [Luo and Zhang, 2018].

The most visible consequence of the noise is the so-called ‘dropout’ effect. Svensson [2020] studied this phenomenon and concluded that the fraction of cells with no count was fully compatible with a noise following a negative binomial distribution. However, this analysis was conducted ignoring the variability in the total number of counts per cell which has a non-negligible effect on the variability of the number of counts for a given gene. In the Supplementary Figure 1 of Svensson [2020], the use of scaled data suggests that the simpler Poisson distribution could be sufficient to explain the number of dropouts.

Here, we reanalyze the same data sets as Svensson [2020], to evaluate all the noise contributions (dropouts and more generally the variability), using scaled data. Following Svensson [2020], we take advantage of published control data sets [Klein et al., 2015; Macosko et al., 2015; Zheng et al., 2017; Svensson et al., 2017] as well as two real biological data sets provided by 10X genomics with cells from a mouse cell line (NIH3T3) and from a human one (HEK293T). In the control data sets, a homogeneous solution of RNA was used as input instead of a solution of single-cells. This allows to study the technical noise without any influence of the cell to cell variability. Indeed, in these data sets, each pseudo single-cell was a droplet of the same RNA solution. On the contrary, the 10X genomics cell lines are real biological data sets where we expect to find both technical variations and biological variations.

When studying gene expression, the quantity of interest is the number of transcripts coding for a given gene *g* in cell *i*. Unfortunately, this quantification is very difficult to obtain and one often focus instead on the proportion of transcripts coding of a given gene *g* out of all transcripts in the cell *i*. We denote by λ_*i,g*_ this fraction. An obvious estimator of λ_*i,g*_ is

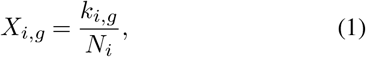

where *k_i,g_* is the number of counts for the gene *g* in the cell *i* and *N_i_* is the total number of counts identified in the cell *i*. However, since *N_i_* is always much smaller than the total number of transcripts in the cell, *k_i,g_* is strongly affected by sampling noise. Rigorously speaking, in the absence of other sources of noise, *k_i,g_* follows a binomial distribution with parameters *N_i_* and λ_i,g_. This distribution can actually be approximated by a Poisson distribution with parameter *N_i_*λ_*i,g*_ for sufficiently small λ_*i,g*_ and large *N_i_*. We thus have

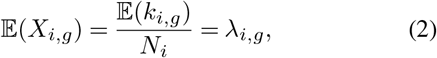

where 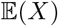 is the expectation of *X*, and

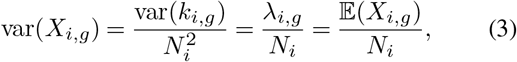

where var(*X*) is the variance of *X*. If additional sources of noise affect the experiment, one would expect an increase in variance. For instance, Svensson [2020] compared the Poisson distribution with the negative binomial distribution with the same expectation and a variance following

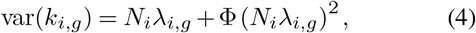

where Φ is a new parameter which allows to account for an additional variance with respect to the Poisson distribution. In particular, the Poisson distribution is recovered for Φ = 0. In terms of *X_i,g_*, the expectation still follows Eq. (2), and the variance is now written as

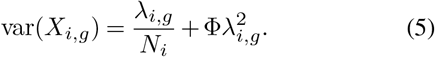

For a given experiment, one only obtains a single realization of *X_i,g_*, so the expectation and variance of *X_i,g_* cannot be measured in order to compare the Poisson and negative binomial distributions. However, in the case of control experiments, each pseudo single cell is sampled from the same RNA solution, so the fraction λ_*i,g*_ is the same in each cell. We denote by λ_*g*_ this common value. By combining the reads in all pseudo cells, one can determine a very precise estimate of λ_*g*_

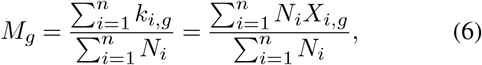

which by construction verifies

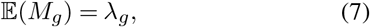

for both the Poisson and negative binomial distributions. We additionally introduce the variance estimator

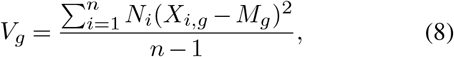

which provides a measure of the spread (variance) of the *X_i,g_* values among the cells for a given gene *g*. The expected value of *V_g_* is

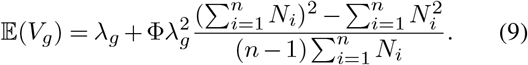

In particular, in the case of the Poisson distribution (Φ = 0), we have 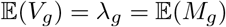, while for Φ > 0 (excess of variance) we obtain 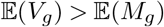.

To estimate which law is more adequate to explain the data we plot *V_g_* against *M_g_* for each gene *g* (Supplementary Fig. 2.1.1). If the data follow the Poisson distribution, we expect the points to align on the *y* = *x* line, while if the data present an additional noise (Φ > 0), the points should end-up above this line. We noticed that the genes corresponding to ERCC (External RNA Control Consortium) spike-in had a behaviour different from what was observed for real genes, especially in the data set from Macosko et al. [2015] where the variance is highly increased (Supplementary Fig. 2.1.1Ab). Thus, we decided to exclude these genes from the analysis. The results with ERCC spike-in information excluded are presented in Fig. 1. Overall, we find a very good agreement of the data with the Poisson distribution in the control data sets (Fig. 1A). In data sets from Svensson et al. [2017] (1) and (2) (Fig. 1Aa and b), the data perfectly follow the Poisson law for all expression levels. In the two other control data sets (from Klein et al. [2015] and Macosko et al. [2015] Fig. 1Ac and d), both models perform equally well until a relatively high level of expression (about 1‰). Above 1‰, the variance slightly deviates from the Poisson prediction. However, in both cases, the deviation is very small and the results at high expression levels suffer from low number statistics. In addition, in the control experiment from Klein et al. [2015], such expression corresponds to the top 0.5% highly expressed genes like ribosomal protein or cytoskeleton proteins. Therefore, we conclude that the Poisson law very well explains the variance observed in the control data sets.

**Table 1.** Comparison of the values *ϕ*_fit_ of the parameter *ϕ* found by Svensson [2020] with the values *ϕ*_Poisson_ expected from the Poisson law when taking into account the variability in the total number of counts per cell (see Eq. (14)).

| Experiment | $\phi_{\text{fit}}$ | $\phi_{\text{Poisson}}$ |
| --- | --- | --- |
| Klein | 0.0428 | 0.0451 |
| Macosko | 0.116 | 0.115 |
| Zheng | 0.0416 | 0.0426 |
| Svensson1 | 0.0940 | 0.0969 |
| Svensson2 | 0.369 | 0.379 |

On the contrary, when considering real experiments, even with a homogeneous single-cell population (Fig. 1B), we find that the variance strongly deviates from the Poisson prediction. Moreover, the deviation increases with the expression level and the negative binomial law captures this behaviour very well. This result clearly demonstrates the inter cellular heterogeneity in so-called homogeneous cell populations.

In Svensson [2020], the author finds that the negative binomial distribution better matches the data than the Poisson law, even for control data sets. This seems to contradict our results. However, the analysis of Svensson [2020] was performed on the raw number of counts *k_i,g_*, without scaling it by the total number of counts *N_i_*. The author uses the following estimators for the mean and variance of *k_i,g_*

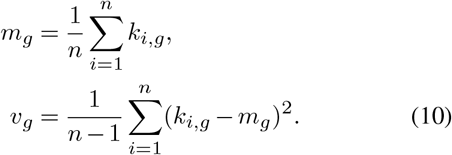

If we assume *k_i,g_* to follow a Poisson distribution with parameter *N_i_*λ_*g*_, the expectation of *m_g_* and *v_g_* are written as

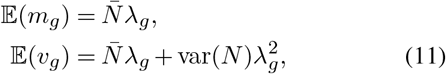

where 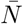 is the average of the *N_i_* values, and var(*N*) their variance

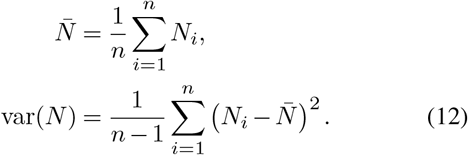

The variations of *N_i_* were neglected by Svensson [2020], which would correspond to assuming 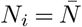 and var(*N*) = 0 in Eq. (11). In this case, one would thus expect to find 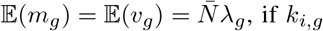, if *k_i,g_* follows a Poisson distribution. Svensson [2020] found the data to be better described by a negative binomial distribution for *k_i,g_* with a free parameters *ϕ* such that

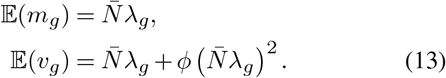

**Fig. 1.**
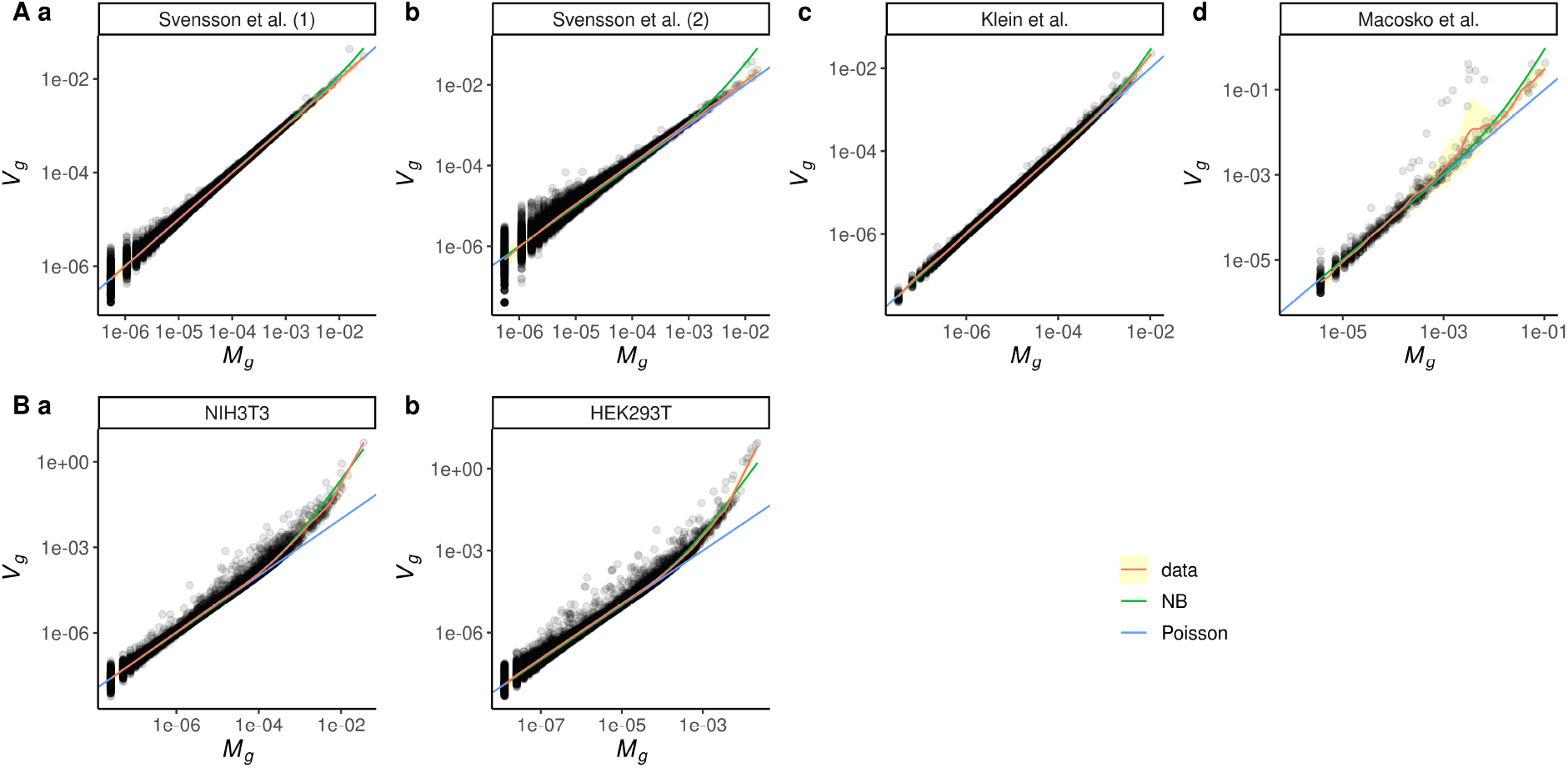
Poisson is a good approximation to explain variance in droplet-based single-cell RNA-seq. In each data set, control data set (**A**) or real single-cell experiment (**B**), the estimator of variance is plotted in function of the estimated mean expression. Each dot is a gene. A Gaussian kernel smoother was applied to evaluate the tendency of the data (red line) and the error around the Gaussian kernel smoothing was estimated (yellow area). The expected variance in the Poisson approximation is a straight line (blue) whereas the expected variance in the negative binomial approximation is a quadratic curve (green). Both axes are in log scale.

This expression can actually match the formula we obtain assuming a Poisson distribution but accounting for the variability of *N_i_* (see Eq. (11)), if we take

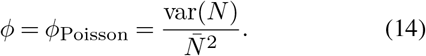

In Table 1, we compare the values of the parameter *ϕ* fitted by Svensson [2020] with the values *ϕ*_Poisson_ computed with Eq. (14), for the control data sets including the spike-in ERCC. We find a very good agreement between these values for all the considered experiments. This shows that the variations in the number of counts per cell is the main source of the excess noise found by Svensson [2020]. This also confirms that the Poisson distribution very well explains the observed variations, as already noticed with the experiment of Fig. 1. Overall, the tests performed in this section show that the variability in droplet-based scRNA-seq is dominated by sampling noise and follows a Poisson distribution. However, it should be noted that we cannot exclude the introduction of biases at specific steps of the technique like increased or decreased reverse transcription or amplification of specific transcripts [Luo and Zhang, 2018]. Indeed such bias would lead to a global increased or decreased presence of these genes and consequently to an overall overestimation or underestimation of each expression value. However, the variance on these genes would still follow a Poisson law.

### Estimating the distribution of expression values of a gene

We are interested here in estimating the intrinsic variability in the expression values of a given gene in a population of cells. As shown above, the output of a droplet-based scRNA-seq experiment presents not only this intrinsic variability but also an additional variability due to sampling noise, following a Poisson distribution. For lowly expressed genes and when the number of counts per cell is low, the sampling noise can actually dominate and hide the intrinsic population variability.

To disentangle these two contributions, we introduce a parametric model of the intrinsic expression variability. We assume that the probability density function (PDF) of the relative expression level of a gene in the population of cells can be approximated by a GMM. The number of components *m* in the mixture, as well as the Gaussians’ amplitudes *A*, means *μ*, and widths *σ*, are free parameters that need to be adjusted to best reproduce the observed number of counts of the considered gene in each cell. We denote by 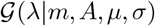 this PDF (see Supplementary 1.1 for more details). Then, for a cell *i* whose expression level for gene *g* is λ_*i,g*_, the probability to get *k_i,g_* out of *N_i_* counts follows the Poisson distribution 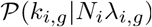. Therefore, the likelihood 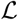 of a given set of parameters (*m, A, μ, σ*) is written as

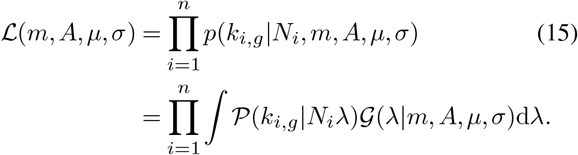

We estimate this integral numerically as described in Supplementary 1.2. While the set of parameters *m, A, μ, σ* could be determined by maximizing the likelihood, we advocate here for a Bayesian approach which prevents an overfit of the data and allows to better estimate confidence intervals on the parameters and on the corresponding PDF G, which is the quantity of interest in this study. We thus use a Markov chain Monte Carlo (MCMC) algorithm to explore the parameters *A, μ, σ* for a given number of components *m*. Then we combine the results obtained for different *m* by evaluating the evidence of each model using an importance sampling algorithm. More details on the algorithms and priors used for this Bayesian approach can be found in Supplementary 1.3 and 1.4. We call this algorithm baredSC for Bayesian Approach to Retrieve Expression Distribution of Single-Cell.

In order to test the efficiency of the algorithm to retrieve the intrinsic distribution, we simulated data using 300, 500, 1561 cells or all (2361 cells) with *N_i_* values taken from a real 10X data set (see Methods). We generated random values for the expression in each cell (λ_*i,g*_) according to various intrinsic distributions: single Gaussian (Fig. 2A), uniform distribution (Fig. 2B), two or three Gaussians (Fig. 2C), or a sub-population of zeros (cells not expressing the gene) and a Gaussian (Fig. 2D). We varied the mean and the width of each of these distributions, to cover a wide range of potential biological cases. Then, we simulated a scRNA-seq by randomly sampling the *k_i,g_* values from a Poisson law using the determined *N_i_* and λ_*i, g*_ values. Finally, these simulated counts were analyzed using baredSC with up to four Gaussians in the mixture.

In the single Gaussian cases, the algorithm very efficiently approximates the intrinsic distribution, even with only 300 cells (see Fig. 2A). We plot in Fig. 2B the case of uniform distributions. While the exact shape of the original distribution cannot be exactly reproduced using a GMM, it is still reasonably well approximated. In particular, when the scale is large enough, the model uses multiple Gaussians to better reproduce a flat distribution. Such a distribution is unlikely to exists in biology, but it illustrates well the versatility of the GMM. When multi-modal distributions were simulated, baredSC could in most cases accurately identify the different modes (see Fig. 2C). The algorithm sometimes slightly deviates from the simulated distribution in the range of low expression when the number of cells is relatively low. This phenomenon is highly linked to the sparsity of the data and to the model’s degeneracy between very low expression levels. Using a higher number of cells allows to break this degeneracy. Finally, in the case of zeros plus a Gaussian, baredSC approximates the distribution with two Gaussians (see Fig. 2D). One of the Gaussians corresponds to the simulated Gaussian. The other is truncated and narrowed close to zero, such that it approximates the sub-population non-expressing cells. However, baredSC tends to predict a lower proportion of expressing cells with a higher mean expression. Again, this phenomenon is linked to the sparsity of the data and to the model’s degeneracy between very low expression and no expression. It disappears when we simulate a higher mean expression for the expressing cells (right column of Fig. 2D). Overall, the algorithm performs very well and largely improves the estimation of the intrinsic expression distribution compared to a classical density plot (see Fig. 2).

### Comparison to Sanity

Sanity [Breda et al., 2021] is a normalization tool aimed at correcting scRNA-seq outputs from sampling noise. The Sanity model shares similar features with baredSC, as both tools use a Bayesian approach to correct from the sampling noise, which is assumed in both cases to follow a Poisson law. However, there are two main differences in the baredSC and Sanity approaches. First, while baredSC uses a GMM to model the intrinsic expression distribution, Sanity uses a single Gaussian model. The Sanity model is thus simpler, which allows for the use of faster algorithms. However, it is also less flexible and precise, especially in cases where the expression distribution is multimodal. Second, Sanity aims at correcting the counts of each cell from the Poisson noise, while baredSC focuses on uncovering the underlying intrinsic expression distribution (PDF) of a gene. The expression PDF can be estimated from Sanity’s outputs by either computing a KDE of the corrected counts, or by computing the posterior distribution from these counts and their error bars [see Breda et al., 2021]. However, these methods are less accurate than the baredSC approach and can degrade the resolution of the PDF.

In order to compare baredSC and Sanity, we generated data using 2361 cells with the same *N_i_* values as for Fig. 2. We generated λ_*i,g*_ values using a single Gaussian, two Gaussians, a single Gaussian in combination with a proportion of cells with no expression, or three Gaussians. In order to more easily compare the results from Sanity and baredSC, the Gaussians were defined in log scale and we run baredSC using the regular log scale. The results are displayed in Fig. 3. For the representation of the normalized counts, the expression of cells with no expression was artificially put to the minimal value. In Fig. 3A where a single Gaussian was simulated, baredSC and the posterior distribution from Sanity overlay with the distribution used to simulate the data. The density from Sanity underestimates the low values and overestimates the mean values. In Fig. 3B, we used a bimodal distribution closed to the one used in the Figure S23 of Breda et al. [2021]. In this case, the posterior distribution from Sanity shows a bi-modal shape, however, while the second Gaussian’s characteristics are well estimated, the first one has a larger scale than expected. Conversely, baredSC finds the characteristics of both Gaussians. In Fig. 3C, where a proportion of cells with no expression was added, baredSC identifies the two sub-populations and gives an inferred PDF very close to the generated one. The posterior distribution from Sanity is composed of a single broad Gaussian missing the bimodality (Gaussian and non-expressing cells). In Fig. 3D, where two Gaussians were used with averages smaller than −7.5, the posterior distribution from Sanity is close to a single Gaussian while baredSC results is close to the expected PDF. Finally, in the last simulation (Fig. 3E) where three Gaussians were used, the posterior distribution from Sanity is very close to the line obtained with normalized counts (Density from data) except that it corrects for cases where there was no detection. This is much less accurate that the PDF provided by baredSC. Overall, in these simulations, baredSC better estimates the distribution compared to Sanity when the distribution is produced by more than a single Gaussian.

These results demonstrate that while Sanity offers a fast solution to correct scRNA-seq outputs from sampling noise, baredSC provides much more accurate results, at the cost of computing time. Both approaches are complementary and have different scopes. The efficiency of the Sanity algorithm allows to apply it massively on all genes. This is especially useful to correct scRNA-seq from sampling noise before applying a clustering and/or a projection algorithm. Such an application would be very intensive in computer time for baredSC, which is dedicated to more in-depth studies of specific genes.

**Fig. 2.**
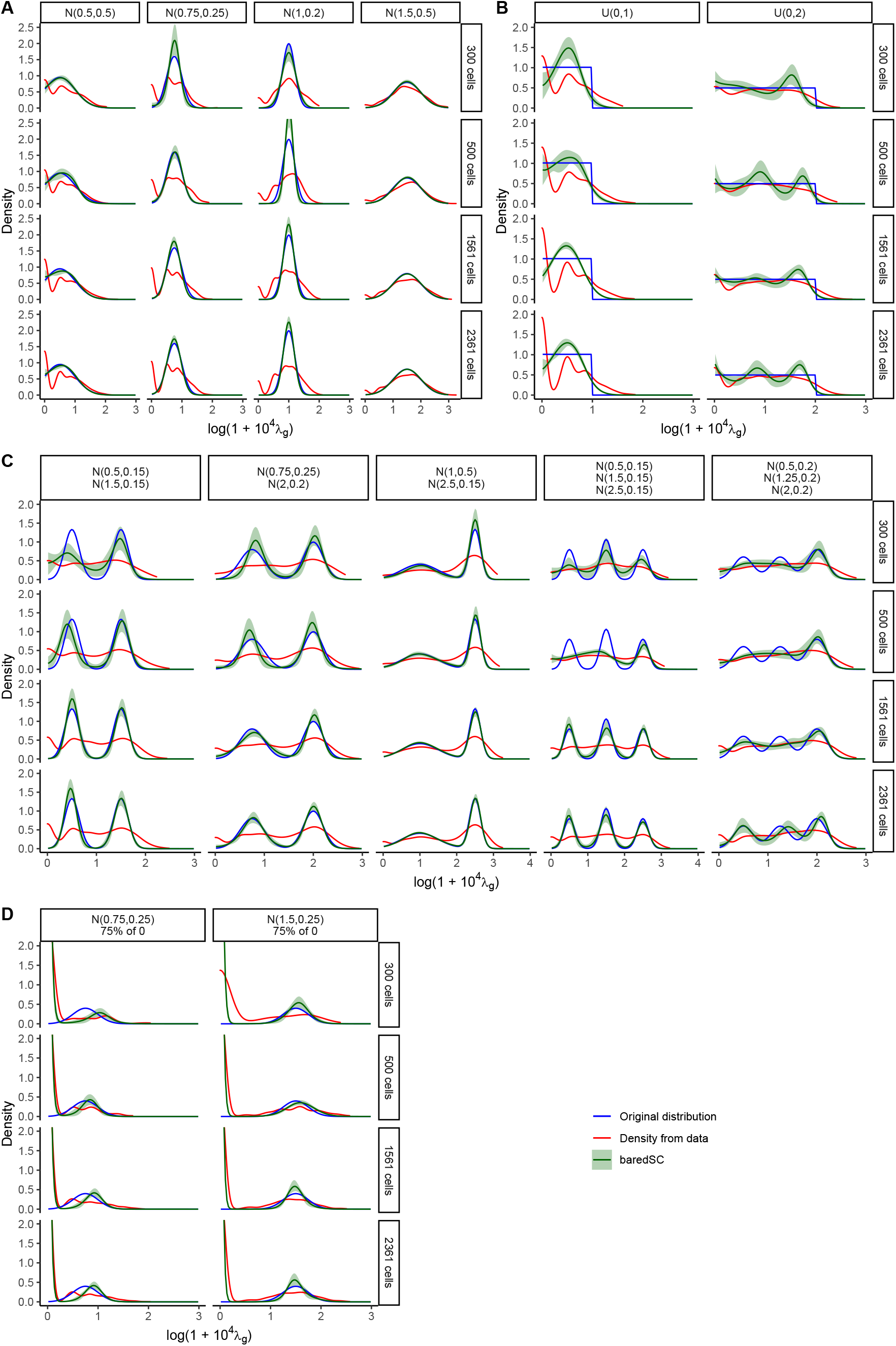
MCMC allows a good estimation of the distribution. In each simulation, the simulated distribution is plotted in blue and its characteristics are written above (N for normal, U for uniform followed by loc and scale values as in the scipy package as well as the proportion of cells with no expression in **D**). The values obtained after Poisson simulation are summarized by the red curve (Density from data). The mean PDF obtained by baredSC is depicted in green and the green area shows the quantile 16-84%. In **A**, distribution is only composed of one normal distribution, in **B**, only one uniform distribution, in **C** two or three Gaussians were used and in **D** a normal distribution for part of the cells and no expression for other.

**Fig. 3.**
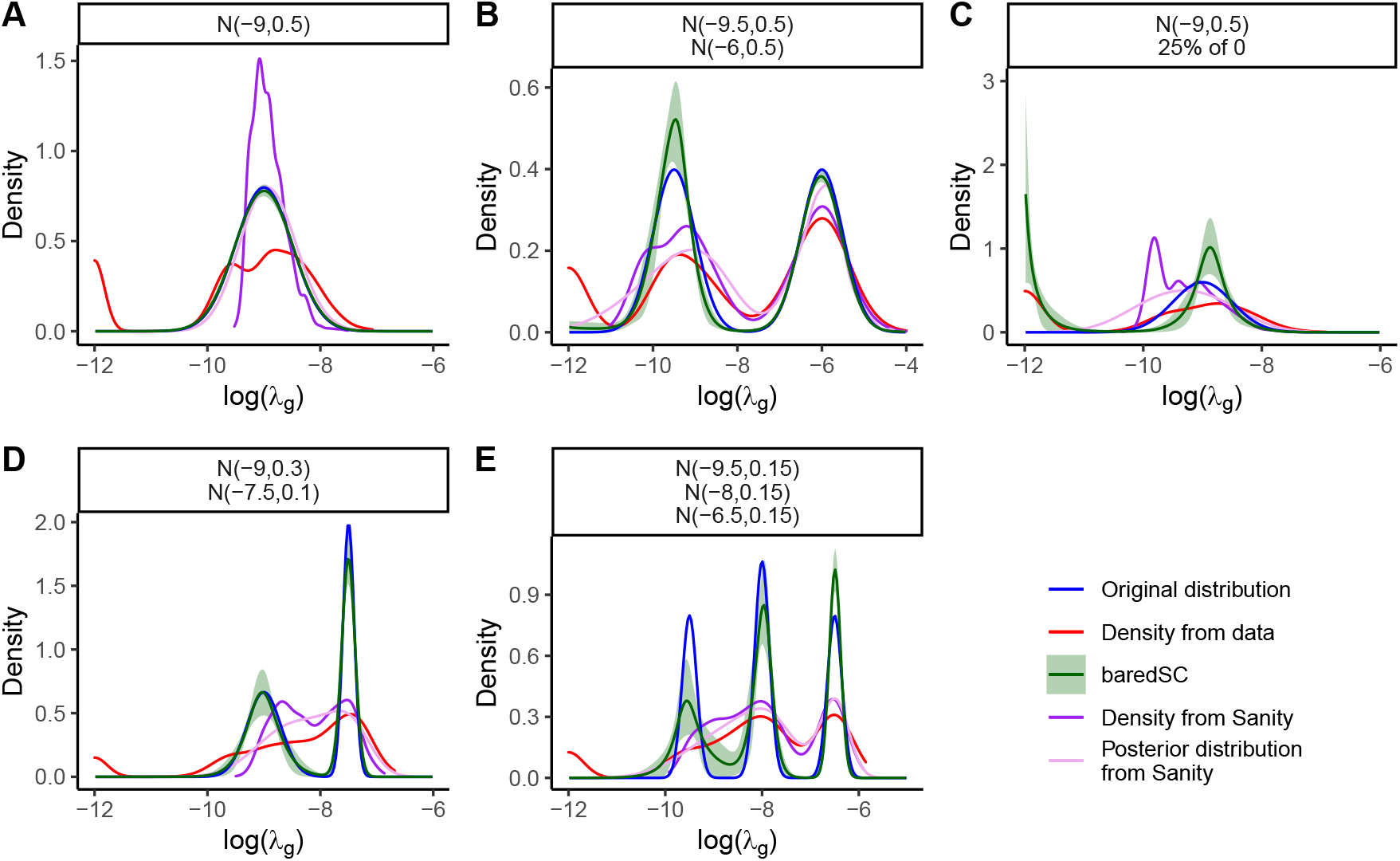
baredSC better estimates the PDF when the distribution is multi-modal. In each simulation, the simulated distribution is plotted in blue and its characteristics are written above (N for normal followed by mean and scale values in log as well as the proportion of cells with no expression). The values obtained after Poisson simulation are summarized by the red curve (Density from data). The mean PDF obtained using baredSC is depicted in green and the green area shows the quantiles at 16-84%. The values obtained using Sanity are summarized by the purple (Density from Sanity) and pink (Posterior distribution from Sanity) curves.

### Estimating the joint distribution of two genes

As shown above, the baredSC approach is able to efficiently recover the expression distribution of a single gene. The same approach can actually be applied to two genes simultaneously to infer the expression distribution in two dimensions (2D). This is of great interest since the study of pairwise correlations between genes can help to better understand gene regulatory networks. Similarly to the case of a single gene, we assume that the distribution can be approximated by a GMM. In the 2D case, each Gaussian in the GMM depends on six parameters: the amplitude, the mean of x and y, the scale of x and y and the correlation. The detailed equations for the 2D application of baredSC can be found in Supplementary 1.5. The same Bayesian approach (MCMC and importance sampling) as in the 1D case is used to explore the parameters. In addition to the parameters and the corresponding expression distribution in 2D, one can deduce from the MCMC posteriors a confidence interval for the Pearson correlation between the expression of the two genes, as well as a one-sided p-value (see Supplementary 1.6).

We simulated data to test the accuracy of baredSC in 2D. We used the same number of cells and the same *N_i_* values as in the 1D case. We generated random values for the expression of both genes x and y in each cell (λ_*i,g*1_ and λ_*i,g*2_) using various distributions described below. Then, we sampled *k*_*i,g*1_ and *k*_*i,g*2_ using Poisson law with parameters λ_*i,g*1_ *N_i_* and λ_*i,g*2_ *N_i_* respectively. These simulated counts were analyzed using baredSC with one to four Gaussians. The results obtained using all cells are presented in Fig. 4, while the results obtained on smaller subgroups are presented in Supplementary Fig. 2.2.2. For each generated data set, we plot in Fig. 4 the distribution used to generate the intrinsic expression values (generated expression), the distribution of normalized simulated raw counts (*k_i,g_/N_i_*, simulated scRNA-seq normalized counts), and the distribution inferred with our approach (baredSC).

We first generated the data using distributions composed of a single truncated 2D Gaussian (Figure 4A). In the three first columns, the mean of *g*1 and *g*2 and the scale of *g*1 and *g*2 are all equal to 0.25. The correlation is set to 0.5 (mainly correlated), −0.5 (mainly anti-correlated) and 0 (independent). We can see that the PDF of the normalized data (simulated scRNA-seq normalized counts) is very noisy and a lot of signal goes on the axes (i.e., one or both genes are not detected). The corresponding Pearson’s correlation coefficient (top left of plot) is close to 0 in the three cases while one would expect values around ±0.3 for the mainly (anti-)correlated cases, in the absence of sampling noise (generated expression). The mean PDF recovered by baredSC (bottom row) is very similar to the original PDF (top row) in these three cases. The Pearson’s correlation coefficient estimated from the MCMC posteriors (top left of each plot in bottom row) is compatible with the value computed from the original PDF (within confidence interval). Moreover, the estimation of the one-sided p-value is significant only in cases where the correlation of the original distribution was non-zero, as expected.

In the three last columns of Fig. 4A, the simulated Gaussians are identical to the first three except that the scale of the gene *g*1 is 0.15 instead of 0.25. Despite the fact that the smaller scale increases the number of drop-out events, baredSC results are still very close to the input PDF.

In Fig. 4B, we split the cells in two equally sized populations. Each population expresses both genes following a 2D Gaussian without correlation. The means of both 2D Gaussians were chosen in order to introduce a correlation (first column of Fig. 4B) or an anti-correlation (second column). The correlation or anti-correlation is already perceptible in the PDF of the normalized counts (simulated scRNA-seq normalized counts). However, the presence of two populations is totally hidden by the sampling noise. The results of baredSC (bottom row) highlight the two distinct populations. The correlation coefficient is very well approximated, especially when compared to the one evaluated on the raw data.

Finally, we generated three different distributions where each of the two genes is expressed in only half of the cells (Fig. 4C). In the first column (each gene expressed in half independent), the cells which do not express the gene *g*1 or the gene *g*2 were chosen independently. In the second column, the two genes are partially anti-correlated. Both genes are expressed in 10 % of the cells, neither is expressed in 10 %, and gene *g*1 (resp. *g*2) is expressed while gene *g*2 (resp. *g*1) is not in 40 %. In the third column, each cell is expressing either *g*1 or *g*2 (each gene expressed in half fully anti-correlated). The estimated correlation coefficients are compatible with the ones obtained on the simulated PDF (generated expression). The shape of the PDF also resembles what was expected. We can however notice that, similarly to what was observed in Fig. 2d for the 1D case, the means of the expressed populations of cells are slightly over estimated. Once again, this can be explained by the sparsity of the data and by the model’s degeneracy.

While it is visually very difficult to interpret the 2D PDF of normalized counts (simulated scRNA-seq normalized counts) for lowly expressed genes, baredSC provides a good approximation of the shape of the original PDF.

### Applications to real data sets

In order to illustrate the power of baredSC and its potential use, we apply it to two data sets.

#### Hoxd13-Hoxa11 anti-correlation in embryonic distal limb

HoxA and HoxD genes are key transcription factors involved in limb patterning [Zakany and Duboule, 2007]. Their expression domain is tightly regulated in space and time. When the limb grows, *Hoxa11* expression has been described as restricted to the proximal domain which will become the arm while *Hoxd13* is expressed in the distal domain which will become the digits. Sheth et al. [2014]; Beccari et al. [2016]; Kherdjemil et al. [2016] showed that *Hoxd13* represses the transcription of *Hoxa11* in the embryonic distal limb leading to two distinct domains of expression. We use the scRNA-seq from Bolt et al. [2021] which has been generated from embryonic forelimbs. A clustering analysis revealed 17 clusters. Using *Hoxd13* as a marker, 4 clusters (3, 4, 5, 9) were attributed to the distal part of the limb [see Bolt et al., 2021]. Unexpectedly, *Hoxa11* was not totally absent from these clusters. We run baredSC on each of these 4 clusters allowing 1 to 4 Gaussians for *Hoxd13* and *Hoxa11* (Fig. 5). In all four clusters, we clearly see a depletion of cells with simultaneous high expression of both genes. As a consequence, the correlation coefficient is significantly negative in all four clusters. These results are thus in agreement with the literature.

#### Multi-modal expression of Pitx1 in embryonic hindlimb

The gene *Pitx1* encodes for a transcription factor expressed in the embryonic hindlimb. It is responsible for the leg patterning. This gene is not expressed in the forelimb and a gain of expression in this domain induces an arm-to-leg phenotype [DeLaurier et al., 2006; Kragesteen et al., 2018]. Its expression is controlled by many enhancers, one of them is *Pen* which accounts for 30-50% of the expression [Kragesteen et al., 2018; Rouco et al., 2021]. Rouco et al. [2021] used scRNA-seq and flow cytometry to investigate whether *Pitx1* was homogeneously expressed across hindlimb cells. They also studied the changes induced by the deletion of the *Pen* enhancer. While scRNA-seq directly measures the level of expression of *Pitx1*, the flow cytometry data measures the level of fluorescence thanks to a GFP-sensor introduced in close proximity to the *Pitx1* promoter. The measure of fluorescence of each cell by flow cytometry should be highly correlated to the level of mRNA of *Pitx1*. However, the exact relationship between the mRNA level and the fluorescence is not known.

In this study, there are three conditions, the forelimb (FL) wild-type (*Pitx1*^+/+^) which has no expression of *Pitx1*, the hindlimb (HL) wild-type which is considered as a tissue expressing *Pitx1* and the HL mutant where the *Pen* enhancer has been deleted (*Pitx1^Pen-/Pen-^*). Both flow cytometry and scRNA-seq data experiments were produced for these 3 samples. We propose a reanalysis of these data. The flow cytometry data are usually represented in log scale (Fig. 6A). It shows the presence of high level of background in absence of expression. Indeed, in the FL *Pitx1*^+/+^ sample in black, the fluorescence level has a relatively wide scale. Rouco et al. [2021] show that the expression of *Pitx1* in the HL wild-type (red curve) is trimodal (Fig. 6A and Fig. 3A of Rouco et al. [2021]). The first mode is included in the range of the FL *Pitx1*^+/+^ so it corresponds to cells which do not express *Pitx1*. Expressing cells can be divided into highly-expressing cells and lowly-expressing cells. The sample HL *Pitx1^Pen-/Pen-^* (blue curve) also exhibits a trimodal distribution but the average expression in low expressing and high expressing cells is decreased compared to the wild-type condition and there is an increased proportion of non expressing cells. These data are consistent with the expected decrease of expression by 30-50%. In order to compare more easily the flow cytometry data with the scRNA-seq expressed in log(1 + 10^4^ *X_i_*), we transformed the fluorescence in log(1 + 0.01 *F_i_*) (Fig. 6B).

**Fig. 4.**
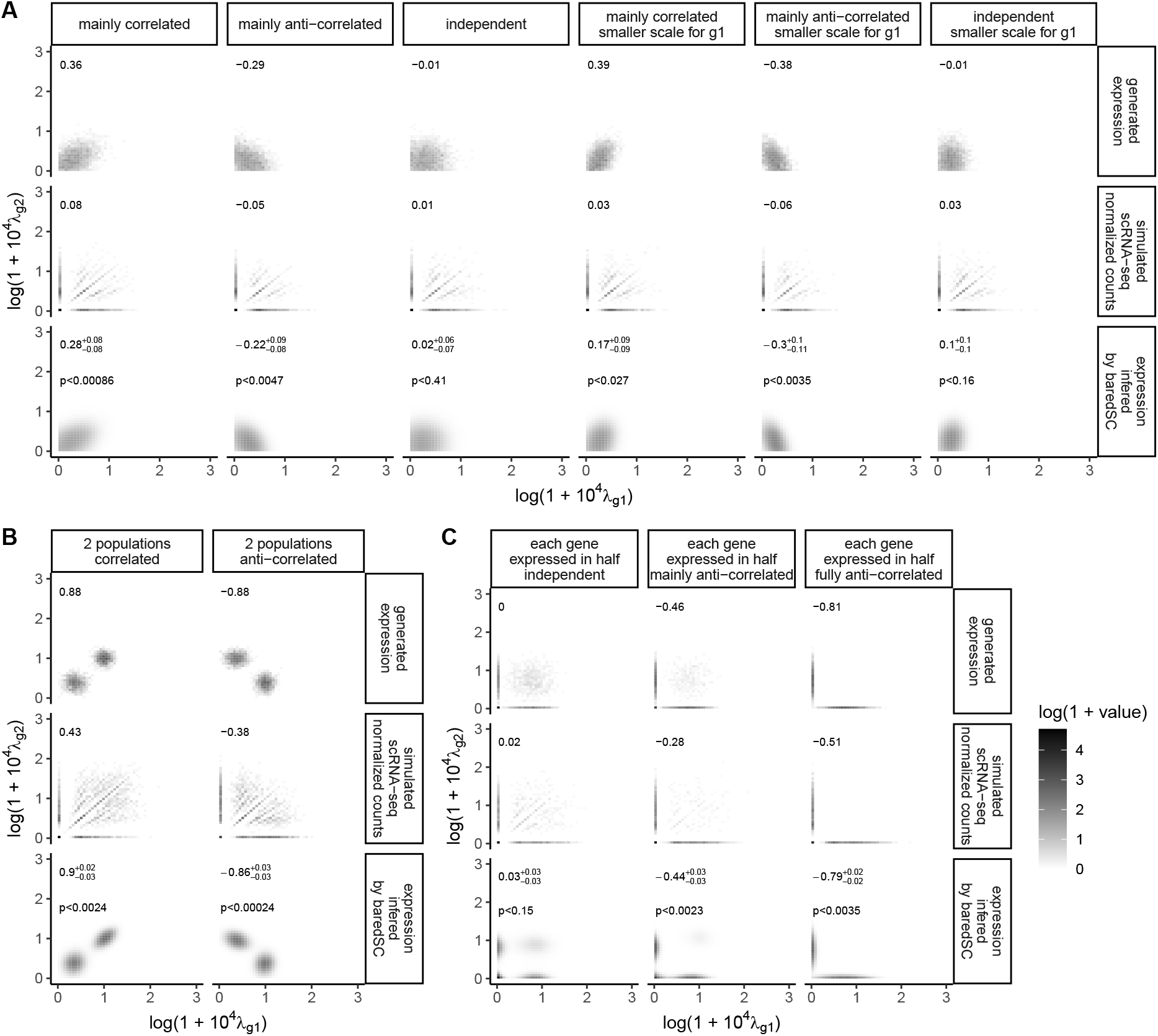
baredSC allows a good estimation of 2D expression distributions. In each simulation, the PDF of the simulated expressions is plotted in the top panel. The values obtained after Poisson simulation are plotted in the middle panel. The mean PDF obtained by baredSC is plotted in the bottom. In the top left corner is written the Pearson’s correlation coefficient and for baredSC its confidence interval. For baredSC, an estimation of the one-sided p-value is displayed. In **A**, distribution is only composed of one truncated normal distribution, in **B**, 2 normal distributions are used, and in **C** each gene is expressed with a Gaussian distribution in half of the cells.

This transformation still highlights the trimodal expression of the HL *Pitx1*^+/+^. When the scRNA-seq results are displayed as density plots (Fig. 6C or Fig. 4A of Rouco et al. [2021]), one cannot distinguish the three modes in the HL *Pitx1*^+/+^. However, classical density plots are very sensitive to sampling noise, and do not allow a precise determination of the expression distribution. Applying baredSC on this data set enables to a better characterization of the shape of the expression distribution (Fig. 6D). The three modes are uncovered: a first one with non-expressing cells approximated by a truncated Gaussian along the y axis, a second one with a mean value around 1, and a third one with a mean value around 2. Because of the background in flow cytometry and the degeneracy of scRNA-seq at very low expression, the comparison of the PDFs from flow cytometry (Fig. 6B) and scRNA-seq (Fig. 6D) on the left part of the plots is difficult. However, at higher expression levels (right part), the PDF are very similar. We also run baredSC in regular log scale (Supplementary Fig. 2.3.3) and confirm this high similarity with the flow cytometry PDF. These results show that the trimodal expression identified by flow cytometry is indeed present in the scRNA-seq. It also demonstrates how baredSC can improve our description of expression variability within a scRNA-seq from a complex tissue.

**Fig. 5.**
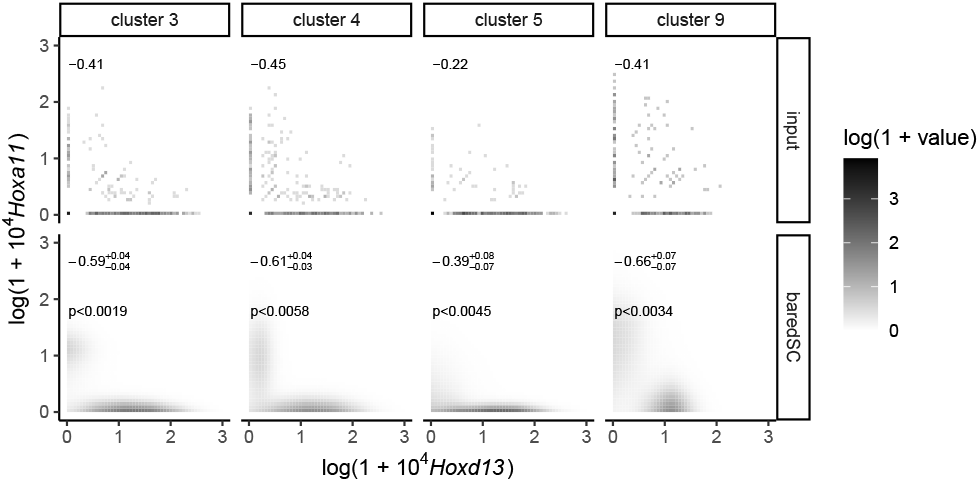
baredSC retrieves the anti-correlation between *Hoxa11* and *Hoxd13* in embryonic distal limb scRNA-seq. Heatmaps showing the PDF of normalized expression of *Hoxa11* and *Hoxd13*. Top panels provide the PDF from input data while bottom panels show the result of baredSC. In the top left corner is written the Pearson’s correlation coefficient and for baredSC its confidence interval. For baredSC, an estimation of the one-sided p-value is displayed.

**Fig. 6.**
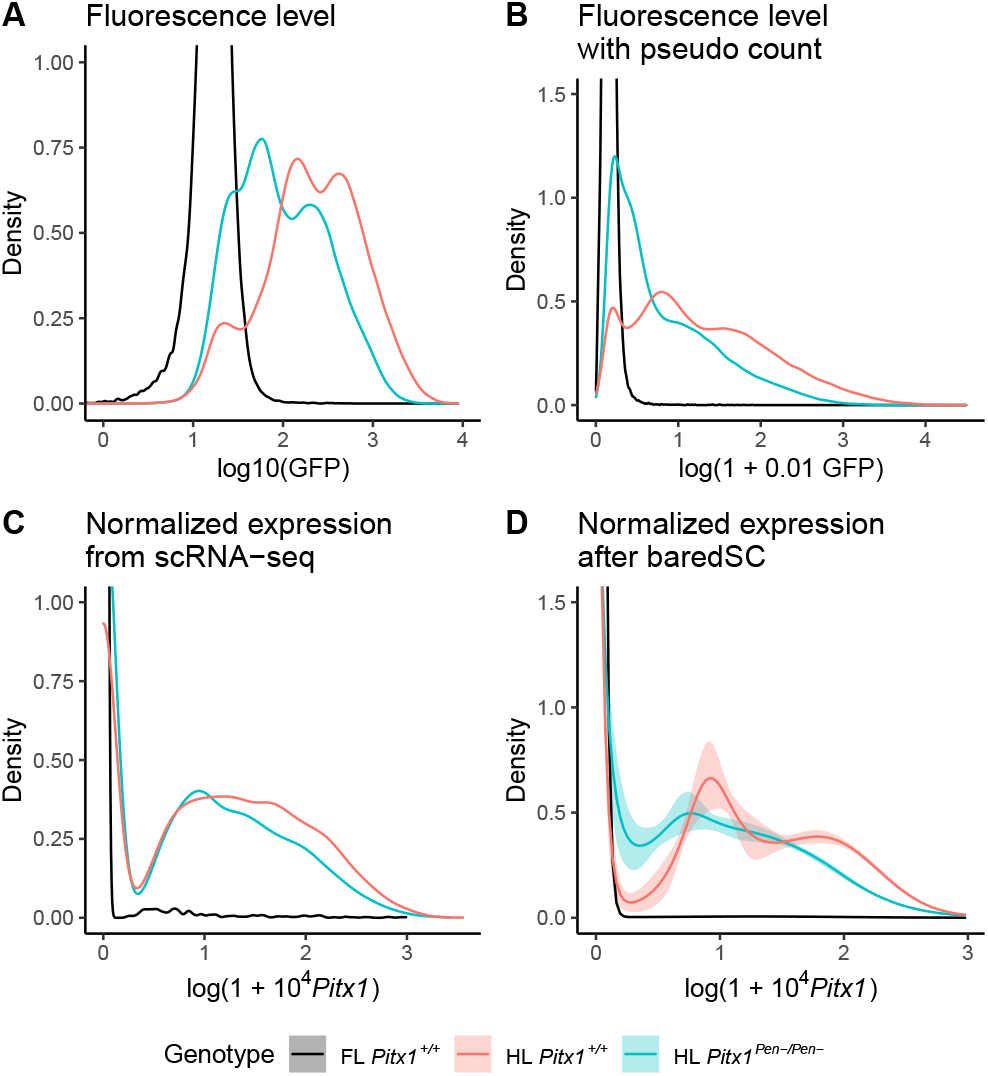
baredSC retrieves the trimodal expression of *Pitx1* in embryonic limb scRNA-seq. **A**: Distribution of fluorescence intensity obtained by flow cytometry in regular log scale. **B**: Distribution of fluorescence intensity on log scale with pseudo count to be closer to the Seurat scale. **C**: Distribution of normalized expression from scRNA-seq as provided by the Seurat package. **D**: Distribution of normalized expression from scRNA-seq as provided by baredSC.

## Conclusion

In this study, we show using control scRNA-seq that most of the technical variability is well approximated by the Poisson distribution. We thus justify its use in mathematical models of droplet-based scRNA-seq. Using simulated data we show that baredSC is a powerful tool to study the distribution of expression levels from droplet-based scRNA-seq of a few genes of interest. It largely outperforms the regularly used density/violin plots to represent the distribution of expression of a gene. It allows to retrieve precisely multi-modal expression distribution even when they are not distinguishable in the input data due to sampling noise. In these cases, baredSC results are more accurate than the posterior distribution from Sanity. The 2D version of baredSC allows better evaluation of the correlation between genes and provides an accurate PDF even for lowly expressed genes.

As it uses a MCMC algorithm to infer the model parameters, the computational cost of the algorithm is non-negligible (typically 10 minutes with 4 CPUs to run 1 to 4 Gaussians, on one of the examples of Fig. 2 using all cells). Additionally, it requires some manual checks, such as the convergence of the MCMC, and the fact that the range of considered expression levels were correctly chosen. This makes this tool more appropriate to in-depth analysis of specific genes than large-scale applications. As shown in the examples, it can be used both on a whole single-cell data set or on cells belonging to specific clusters. Potential applications for a single gene are modality description or estimation of proportion of cells expressing a given gene as well as comparisons between samples or clusters. We apply it to a real biological scRNA-seq data set to study the modality of the expression of *Pitx1*. We demonstrate that the fluorescence measurement of the GFP sensor, integrated in close proximity of *Pitx1* promoter, is highly correlated with the mRNA level of the gene. The inference of PDF in 2D is of great interest for the study of genes’ interactions or its comparison between clusters of cell. We apply it to confirm the anti-correlation between *Hoxa11* and *Hoxd13* in the developing limb.

## Method

Most of the details regarding the equations and baredSC implementation are in the Supplementary Material.

### Software and code availability

baredSC is a python opensource package hosted on github at https://github.com/lldelisle/baredSC and can be installed by pip. For this paper, version 1.0.0 was used with default parameters. The number of samples, starting at 100 000 was increased by 10 times until reaching a number of effective samples above 200.

All the scripts needed to reproduce all the figures of the paper are available at https://github.com/lldelisle/scriptsForLopezDelisleEtAl2021. The figures were made in R (https://www.r-project.org/) with ggplot2 [Wickham, 2016] and ggpubr (https://rpkgs.datanovia.com/ggpubr/).

### Simulation of data

In order to get realistic values for *N_i_*, we extracted the *N_i_* from the cells which were attributed to NIH3T3 in the data set provided by 10X with the same criteria used by [Svensson, 2020]. Then, 300 cells were randomly attributed to a first group, 500 cells to another one and the others to a third one. The expressions were then generated by python scripts available at https://github.com/lldelisle/scriptsForLopezDelisleEtAl2021.

## AUTHOR CONTRIBUTIONS

L. Lopez-Delisle wrapped the software to implement different options, simulated data and ran the software, wrote the manuscript.

J.-B. Delisle coded the core of the software, brought his expertise in statistics, wrote the manuscript.

## ACKNOWLEDGEMENTS

We thank all members of the D. Duboule and G. Andrey laboratories for their inputs. We acknowledge particularly C.C. Bolt for providing the scRNA-seq data for the *Hoxd13-Hoxa11* application and R. Rouco and O. Bompadre for sharing the flow cytometry and the scRNA-seq data for the *Pitx1* application. L.L-D was supported by a grant from the European Research Council (RegulHox, #588029, to D.D.) and by EPFL. J.-B. D. acknowledges support by the Swiss National Science Foundation (SNSF). This work has, in part, been carried out within the framework of the National Centre for Competence in Research PlanetS supported by SNSF.

## Bibliography

Bacher, R. and Kendziorski, C. (2016). Design and computational analysis of single-cell RNA-sequencing experiments. Genome Biology, 17(1):63.

Bartel, D. P. (2018). Metazoan MicroRNAs. Cell, 173(1):20–51.

Beccari, L., Yakushiji-Kaminatsui, N., Woltering, J. M., Necsulea, A., Lonfat, N., Rodríguez-Carballo, E., Mascrez, B., Yamamoto, S., Kuroiwa, A., and Duboule, D. (2016). A role for HOX13 proteins in the regulatory switch between TADs at the HoxD locus. Genes & Development, 30(10):1172–1186.

Bolt, C. C., Lopez-Delisle, L., Mascrez, B., and Duboule, D. (2021). MESOMELIC DYSPLASIAS ASSOCIATED WITH THE HOXD LOCUS ARE CAUSED BY REGULATORY REALLOCATIONS. bioRxiv, page 2021.02.01.429171. Publisher: Cold Spring Harbor Laboratory Section: New Results.

Breda, J., Zavolan, M., and van Nimwegen, E. (2021). Bayesian inference of gene expression states from single-cell RNA-seq data. Nature Biotechnology, pages 1–9. Publisher: Nature Publishing Group.

Butler, A., Hoffman, P., Smibert, P., Papalexi, E., and Satija, R. (2018). Integrating single-cell transcriptomic data across different conditions, technologies, and species. Nature Biotechnology, 36(5):411–420. Number: 5 Publisher: Nature Publishing Group.

DeLaurier, A., Schweitzer, R., and Logan, M. (2006). Pitx1 determines the morphology of muscle, tendon, and bones of the hindlimb. Developmental Biology, 299(1):22–34.

Haque, A., Engel, J., Teichmann, S. A., and Lönnberg, T. (2017). A practical guide to singlecell RNA-sequencing for biomedical research and clinical applications. Genome Medicine, 9(1):75.

Kherdjemil, Y., Lalonde, R. L., Sheth, R., Dumouchel, A., de Martino, G., Pineault, K. M., Wellik, D. M., Stadler, H. S., Akimenko, M.-A., and Kmita, M. (2016). Evolution of Hoxa11 regulation in vertebrates is linked to the pentadactyl state. Nature, 539(7627):89–92.

Klein, A. M., Mazutis, L., Akartuna, I., Tallapragada, N., Veres, A., Li, V., Peshkin, L., Weitz, D. A., and Kirschner, M. W. (2015). Droplet barcoding for single-cell transcriptomics applied to embryonic stem cells. Cell, 161(5):1187–1201.

Kragesteen, B. K., Spielmann, M., Paliou, C., Heinrich, V., Schöpflin, R., Esposito, A., Annunziatella, C., Bianco, S., Chiariello, A. M., Jerković, I., Harabula, I., Guckelberger, P., Pechstein, M., Wittler, L., Chan, W.-L., Franke, M., Lupiáñez, D. G., Kraft, K., Timmermann, B., Vingron, M., Visel, A., Nicodemi, M., Mundlos, S., and Andrey, G. (2018). Dynamic 3D chromatin architecture contributes to enhancer specificity and limb morphogenesis. Nature Genetics, 50(10):1463–1473.

Luo, Q. and Zhang, H. (2018). Emergence of Bias During the Synthesis and Amplification of cDNA for scRNA-seq. Advances in Experimental Medicine and Biology, 1068:149–158.

Macosko, E. Z., Basu, A., Satija, R., Nemesh, J., Shekhar, K., Goldman, M., Tirosh, I., Bialas, A. R., Kamitaki, N., Martersteck, E. M., Trombetta, J. J., Weitz, D. A., Sanes, J. R., Shalek, A. K., Regev, A., and McCarroll, S. A. (2015). Highly Parallel Genome-wide Expression Profiling of Individual Cells Using Nanoliter Droplets. Cell, 161(5):1202–1214.

Nayak, R. and Hasija, Y. (2021). A hitchhiker’s guide to single-cell transcriptomics and data analysis pipelines. Genomics, 113(2):606–619.

Pollen, A. A., Nowakowski, T. J., Shuga, J., Wang, X., Leyrat, A. A., Lui, J. H., Li, N., Szpankowski, L., Fowler, B., Chen, P., Ramalingam, N., Sun, G., Thu, M., Norris, M., Lebofsky, R., Toppani, D., Kemp, D. W., Wong, M., Clerkson, B., Jones, B. N., Wu, S., Knutsson, L., Alvarado, B., Wang, J., Weaver, L. S., May, A. P., Jones, R. C., Unger, M. A., Kriegstein, A. R., and West, J. A. A. (2014). Low-coverage single-cell mRNA sequencing reveals cellular heterogeneity and activated signaling pathways in developing cerebral cortex. Nature Biotechnology, 32(10):1053–1058.

Rouco, R., Bompadre, O., Rauseo, A., Fazio, O., Thorel, F., Peraldi, R., and Andrey, G. (2021). Cell-specific alterations in Pitx1 regulatory landscape activation caused by the loss of a single enhancer. bioRxiv, page 2021.03.10.434611. Publisher: Cold Spring Harbor Laboratory Section: New Results.

Segal, E., Shapira, M., Regev, A., Pe’er, D., Botstein, D., Koller, D., and Friedman, N. (2003). Module networks: identifying regulatory modules and their condition-specific regulators from gene expression data. Nature Genetics, 34(2):166–176. Number: 2 Publisher: Nature Publishing Group.

Sheth, R., Bastida, M. F., Kmita, M., and Ros, M. (2014). “Self-regulation,” a new facet of Hox genes’ function. Developmental Dynamics: An Official Publication of the American Association of Anatomists, 243(1):182–191.

Svensson, V. (2020). Droplet scRNA-seq is not zero-inflated. Nature Biotechnology, 38(2):147–150. Number: 2 Publisher: Nature Publishing Group.

Svensson, V., Natarajan, K. N., Ly, L.-H., Miragaia, R. J., Labalette, C., Macaulay, I. C., Cvejic, A., and Teichmann, S. A. (2017). Power analysis of single-cell RNA-sequencing experiments. Nature Methods, 14(4):381–387.

Tanay, A. and Regev, A. (2017). Scaling single-cell genomics from phenomenology to mechanism. Nature, 541(7637):331–338. Number: 7637 Publisher: Nature Publishing Group.

Tang, F., Barbacioru, C., Wang, Y., Nordman, E., Lee, C., Xu, N., Wang, X., Bodeau, J., Tuch, B. B., Siddiqui, A., Lao, K., and Surani, M. A. (2009). mRNA-Seq whole-transcriptome analysis of a single cell. Nature Methods, 6(5):377–382.

Tarbier, M., Mackowiak, S. D., Frade, J., Catuara-Solarz, S., Biryukova, I., Gelali, E., Menéndez, D. B., Zapata, L., Ossowski, S., Bienko, M., Gallant, C. J., and Friedländer, M. R. (2020). Nuclear gene proximity and protein interactions shape transcript covariations in mammalian single cells. Nature Communications, 11(1):5445. Number: 1 Publisher: Nature Publishing Group.

Vallejos, C. A., Risso, D., Scialdone, A., Dudoit, S., and Marioni, J. C. (2017). Normalizing singlecell RNA sequencing data: challenges and opportunities. Nature Methods, 14(6):565–571. Number: 6 Publisher: Nature Publishing Group.

Wang, T., Li, B., Nelson, C. E., and Nabavi, S. (2019). Comparative analysis of differential gene expression analysis tools for single-cell RNA sequencing data. BMC Bioinformatics, 20(1):40.

Wickham, H. (2016). ggplot2: Elegant Graphics for Data Analysis. Springer-Verlag New York.

Wolf, F. A., Angerer, P., and Theis, F. J. (2018). SCANPY: large-scale single-cell gene expression data analysis. Genome Biology, 19(1):15.

Zakany, J. and Duboule, D. (2007). The role of Hox genes during vertebrate limb development. Current Opinion in Genetics & Development, 17(4):359–366.

Zheng, G. X. Y., Terry, J. M., Belgrader, P., Ryvkin, P., Bent, Z. W., Wilson, R., Ziraldo, S. B., Wheeler, T. D., McDermott, G. P., Zhu, J., Gregory, M. T., Shuga, J., Montesclaros, L., Underwood, J. G., Masquelier, D. A., Nishimura, S. Y., Schnall-Levin, M., Wyatt, P. W., Hindson, C. M., Bharadwaj, R., Wong, A., Ness, K. D., Beppu, L. W., Deeg, H. J., McFarland, C., Loeb, K. R., Valente, W. J., Ericson, N. G., Stevens, E. A., Radich, J. P., Mikkelsen, T. S., Hindson, B. J., and Bielas, J. H. (2017). Massively parallel digital transcriptional profiling of single cells. Nature Communications, 8:14049.

