## Supplementary Material for "baredSC: Bayesian Approach to Retrieve Expression Distribution of Single-Cell"

May 26, 2021

#### 1 Supplementary methods

##### 1.1 GMM of the PDF

In the scRNA-seq field, the most popular analysis packages (Seurat and Scanpy) uses a log transformation of the normalized counts to be able to see both highly expressing cells and lowly expressing cells. To handle the numerous cases where the gene was not detected, they use a transformation of type $log(1 + N_t X_{i,g})$ where $N_t$ is the targeted number of counts used to normalize $X_{i,g}$ values. In the Seurat package $N_t$ is fixed to $10^4$ while in Scanpy it is set by default to the median of $N_i$ or it can be user defined. However, the pseudo-count in log may introduce a distortion and users may prefer to work in regular log scale. In baredSC, both scales have been implemented, and the PDF is modeled by a fixed number of Gaussians in the chosen scale. We define

$$x_g = \log(1 + N_t \lambda_g), \tag{1}$$

in the case of the Seurat/Scanpy scale or

$$x_g = \log(\lambda_g), \tag{2}$$

in the regular log scale case. We denote by $l$ the mapping between $x$ and $\lambda$: $x = l(\lambda)$ and $\lambda = l^{-1}(x)$. The likelihood is:

$$\begin{aligned}
\mathcal{L}(\theta) &= \prod_{i=1}^{n} p(k_{i,g}|N_i, \theta) \qquad (3)\\
&= \prod_{i=1}^{n} \int_{\lambda_{\min}}^{\lambda_{\max}} \mathcal{P}(k_{i,g}|N_i\lambda)\mathcal{G}(\lambda|\theta)\mathrm{d}\lambda\\
&= \prod_{i=1}^{n} \int_{x_{\min}}^{x_{\max}} \mathcal{P}(k_{i,g}|N_i l^{-1}(x))G(x|\theta)\mathrm{d}x, \qquad (4)
\end{aligned}$$

where $\theta = (m, A, \mu, \sigma)$ is the set of model parameters, and

$$
\begin{aligned}
G(x|\theta) &= \sum_{j=1}^{m} a_j \exp\left(-\frac{(x-\mu_j)^2}{2\sigma_j^2}\right), \\
a_j &= A_j \left(\int_{x_{\min}}^{x_{\max}} \exp\left(-\frac{(x-\mu_j)^2}{2\sigma_j^2}\right) \mathrm{d}x\right)^{-1}, \\
\mathcal{G}(\lambda|\theta) &= G(l(\lambda)|\theta) l'(\lambda).
\end{aligned} \tag{5}
$$

#### 1.2 Estimation of likelihood

Unfortunately, the integral in Eq. (4) cannot be computed analytically, and we thus evaluate it numerically using a Riemann sum

$$
\begin{aligned}
L_i(\theta) &= \int_{x_{\min}}^{x_{\max}} \mathcal{P}(k_{i,g}|N_i l^{-1}(x)) G(x|\theta) \mathrm{d}x \\
&\approx \sum_{q=1}^{n_x} \mathcal{P}(k_{i,g}|N_i l^{-1}(x_q)) G(x_q|\theta) \delta x,
\end{aligned} \tag{6}
$$

where $x_q$ are the centers of the $n_x$ equally sized bins, and $\delta x = (x_{\max} - x_{\min})/n_x$ is the width of each bin. In this integral, the Poisson part $(\mathcal{P}(k_{i,g}|N_i l^{-1}(x)))$ does not depend on the model parameters. It can thus be pre-computed on a given grid of $x$ values for each cell, and stored in a $n \times n_x$ matrix $P$. On the contrary, the GMM part $(G(x|\theta))$ depends on the model parameters but is the same for all cells. Therefore, for a given set of parameters, it must be computed on the grid of $x$ only once for all cells, and stored in a vector $\gamma(\theta)$ of size $n_x$. We thus obtain the vector $L$ as the dot product

$$
L(\theta) \approx P\gamma(\theta). \tag{7}
$$

Then the likelihood is simply given by

$$
\mathcal{L}(\theta) = \prod_{i=1}^{n} L_i(\theta). \tag{8}
$$

The computational cost of this method is dominated by the dot product of Eq. (7). Indeed, the matrix $P$ can be pre-computed so its cost is negligible, the cost of computing the vector $\gamma$ scales as $\mathcal{O}(n_x)$, while the cost of the dot product scales as $\mathcal{O}(nn_x)$. Therefore, a trade-off between resolution and cost must be found to choose the number of bins $n_x$.

However, since the cost of computing $P$ and $\gamma$ is much lower than the cost of the dot product, $P$ and $\gamma$ can actually be computed on a finer grid to improve precision. We thus subdivide each of the $n_x$ original bins into $n_s$ sub-bins and compute the average value of $\mathcal{P}$ over these sub-bins. Similarly, each of the $n_x$ original bins were divided into $n_r$ sub-bins and compute the average value of $G$ over these sub-bins.

$$
\begin{aligned}
P_{i,q} &= \frac{1}{n_s} \sum_{s=1}^{n_s} \mathcal{P}(k_{i,g}|N_i l^{-1}(x_{q,s})), \\
\gamma_q(\theta) &= \frac{1}{n_r} \sum_{r=1}^{n_r} G(x_{q,r}|\theta).
\end{aligned} \tag{9}
$$

#### 1.3 Priors on parameters

We chose the priors on the GMM parameters as follows

- number of Gaussians $m$: uniform distribution over $[\![1, m_{\max}]\!]$ (with $m_{\max} = 4$ in this article);
- amplitudes $A_j$: uniform distribution over $[0, 1]$, with the additional condition $\sum_{j=1}^{m} A_j = 1$;
- means $\mu_j$: uniform distribution over $[x_{\min} - 3\sigma_j, x_{\max} + 3\sigma_j]$;
- scales $\sigma_j$: log-uniform distribution over $[\sigma_{\min}, x_{\max} - x_{\min}]$ (with $\sigma_{\min} = 0.1$ in this article).

This model is symmetric in the sense that two Gaussians can be arbitrarily swapped without changing the results. This degeneracy can deteriorate the convergence of the MCMC algorithm. We thus prevent any swapping to ensure the uniqueness of the solution.

#### 1.4 MCMC algorithm

We use the samsam python package (`https://gitlab.unige.ch/Jean-Baptiste.Delisle/samsam`) which is a scaled adaptive Metropolis algorithm [Haario et al., 2001; Andrieu and Thoms, 2008; Delisle et al., 2018]. We first apply a burning phase with simulated annealing in order to prevent the MCMC from being trapped in local minima. After this burning phase, the proper MCMC is run to sample the posterior distribution.

#### 1.5 baredSC in 2D

The extension of baredSC to the 2D case is very similar to the 1D case. We use either the "Seurat" scale (Eq. (1)) or the regular log scale (Eq. (2)) and define $x_1 = l(\lambda_{g_1})$, $x_2 = l(\lambda_{g_2})$ for two genes $g_1$ and $g_2$, where $l$ is the chosen scale mapping. Each Gaussian of the GMM is defined by 6 parameters: the amplitude ($A_j$), the means of $x_1$ and $x_2$ ($\mu_{1,j}$ and $\mu_{2,j}$), their scales ($\sigma_{1,j}$ and $\sigma_{2,j}$), and the correlation ($\rho_j$). The likelihood is written as

$$\begin{aligned}\mathcal{L}(\Theta) &= \prod_{i=1}^{n} p(k_{i,g}|N_i, \Theta)\\ &= \prod_{i=1}^{n} \int_{x_{2,\min}}^{x_{2,\max}} \int_{x_{1,\min}}^{x_{1,\max}} \mathcal{P}(k_{i,g_1}|N_i l^{-1}(x_1))\mathcal{P}(k_{i,g_2}|N_i l^{-1}(x_2))G(x|\Theta)\mathrm{d}x,\end{aligned} \tag{10}$$

where $\Theta = (m, A, \mu_1, \mu_2, \sigma_1, \sigma_2, \rho)$ is the set of model parameters, and

$$\begin{aligned}G(x|\Theta) &= \sum_{j=1}^{m} a_j \exp\left(-\frac{1}{2}(x - \mu_{.,j})^{\mathrm{T}} C_j^{-1}(x - \mu_{.,j})\right),\\ a_j &= A_j \left(\int_{x_{2,\min}}^{x_{2,\max}} \int_{x_{1,\min}}^{x_{1,\max}} \exp\left(-\frac{1}{2}(x - \mu_{.,j})^{\mathrm{T}} C_j^{-1}(x - \mu_{.,j})\right)\mathrm{d}x\right)^{-1},\end{aligned} \tag{11}$$

with

$$C_j = \begin{pmatrix} \sigma_{1,j}^2 & \sigma_{1,j}\sigma_{2,j}\rho_j \\ \sigma_{1,j}\sigma_{2,j}\rho_j & \sigma_{2,j}^2 \end{pmatrix}. \tag{12}$$

The double integral in Eq. (10) is computed numerically in a very similar manner as in the 1D case, but this time by defining bins on a 2D grid. The priors are the same as in the 1D case, and we take a prior for the correlation $\rho_j$ with a normal distribution of mean 0 and scale 0.3 truncated over $[-0.95, 0.95]$ in order to limit the false positive (anti-)correlation detection.

#### 1.6 Estimation of the p-value for the correlation

If the correlation between the expression of two genes is suspected to be of a given sign $s$, we define the one-sided p-value $\alpha$ as the probability for the correlation to actually be of the opposite sign $-s$. To estimate this p-value, we first compute the Pearson's correlation coefficient from the 2D PDF for each sample of the MCMC. We thus obtain samples from the posterior distribution of the correlation coefficient. Let's denote by $k$ the number of independent samples for which the correlation is of sign $-s$ and $n$ the total number of independent samples. We have (Bayes formula)

$$p(\alpha|k) = \frac{p(k|\alpha)p(\alpha)}{p(k)}, \tag{13}$$

where $p(k|\alpha)$ can be approximated by a Poisson distribution of parameter $n\alpha$ (in the limit of low $\alpha$ and high $n$)

$$p(k|\alpha) = \frac{(n\alpha)^k}{k!} e^{-n\alpha}. \tag{14}$$

We additionally assume a uniform prior for $\alpha$. The conditional expectation of $\alpha$ knowing $k$ is thus

$$\mathbb{E}(\alpha|k) = \int_0^1 \alpha p(\alpha|k) \mathrm{d}\alpha = \frac{k+1}{n} \frac{I_{k+1}}{I_k}, \tag{15}$$

with

$$I_k = \int_0^1 \frac{(n\alpha)^k e^{-n\alpha}}{k!} \mathrm{d}\alpha = \frac{1}{n} \left( 1 - e^{-n} \sum_{j=0}^{k} \frac{n^j}{j!} \right). \tag{16}$$

For $n$ sufficiently large, we have

$$I_k \approx \frac{1}{n}, \tag{17}$$

and

$$\mathbb{E}(\alpha|k) \approx \frac{k+1}{n}. \tag{18}$$

Similarly one can show that the variance is $\mathrm{var}(\alpha|k) \approx \frac{k+1}{n^2}$. The p-value provided on Fig. 4 and Fig. 5 of the article and Supplementary Fig. 2.2.2 is the 1-$\sigma$ upper limit $p < \mathbb{E}(\alpha|k) + \sqrt{\mathrm{var}(\alpha|k)} = \frac{k+1+\sqrt{k+1}}{n}$.

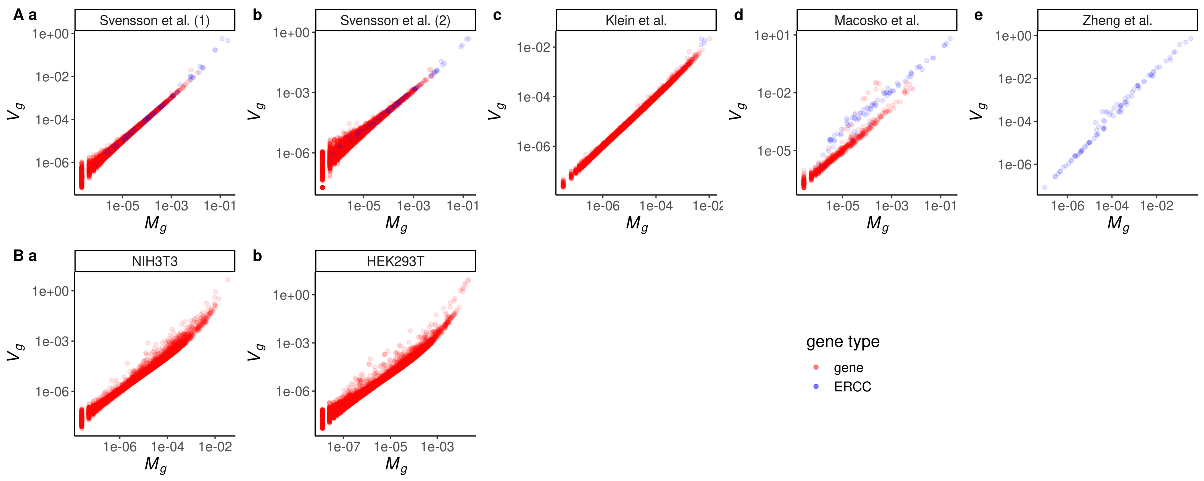


**Supplementary Fig. 2.1.1: ERCC exhibit an increased variance compare to mRNA coding for genes.** In each data set, control data set (**A**) or real single-cell experiment (**B**), the estimator of variance is plotted as a function of the estimated mean expression. Each dot is a gene. The dots are colored by gene type, spike-in ERCC (External RNA Control Consortium) are depicted in blue while mRNA are depicted in red.

### 2 Supplementary results

#### 2.1 ERCC show an increase variance

When scRNA-seq experiments are performed, some manufacturers recommend to spike-in ERCC (External RNA Control Consortium). These are synthetic RNA molecules of various length and various GC contents which can be added to the RNA preparation and sequenced along with the mRNA. They can thus be used as controls to normalize the data. The control experiments that we analyze here all include ERCC. In Supplementary Fig. 2.1.1, we plot the variance as a function of average expression for real and ERCC genes. We notice that the ERCC genes have an increased variance, and are sometimes expressed at a much higher level than regular genes. In order to avoid contaminating our results with this peculiar behavior of ERCC genes, we exclude them from our analysis.

#### 2.2 baredSC in 2 dimensions performs well even with smaller number of cells

While in the main figure, all simulations were run for the whole set of cells (2361), we also run baredSC when only a smaller number of cells was provided (Supplementary Fig. 2.2.2). With 1561 cells, the results are very close to the results obtained for the 2361 cells. The only exception is the case of each gene expressed in half independent (Supplementary Fig. 2.2.2C) where it detects a significant but very modest correlation (0.09). For the smaller numbers of cells (300 and 500), it globally performs very well especially in simulations with multi-modal distributions. In Supplementary Fig. 2.2.2A where the expression is very low, the algorithm fails to detect most of the (anti-)correlations but the average PDF is close to the simulated one. In this range of expression, a high number of cell is necessary to infer a good correlation of the distribution.

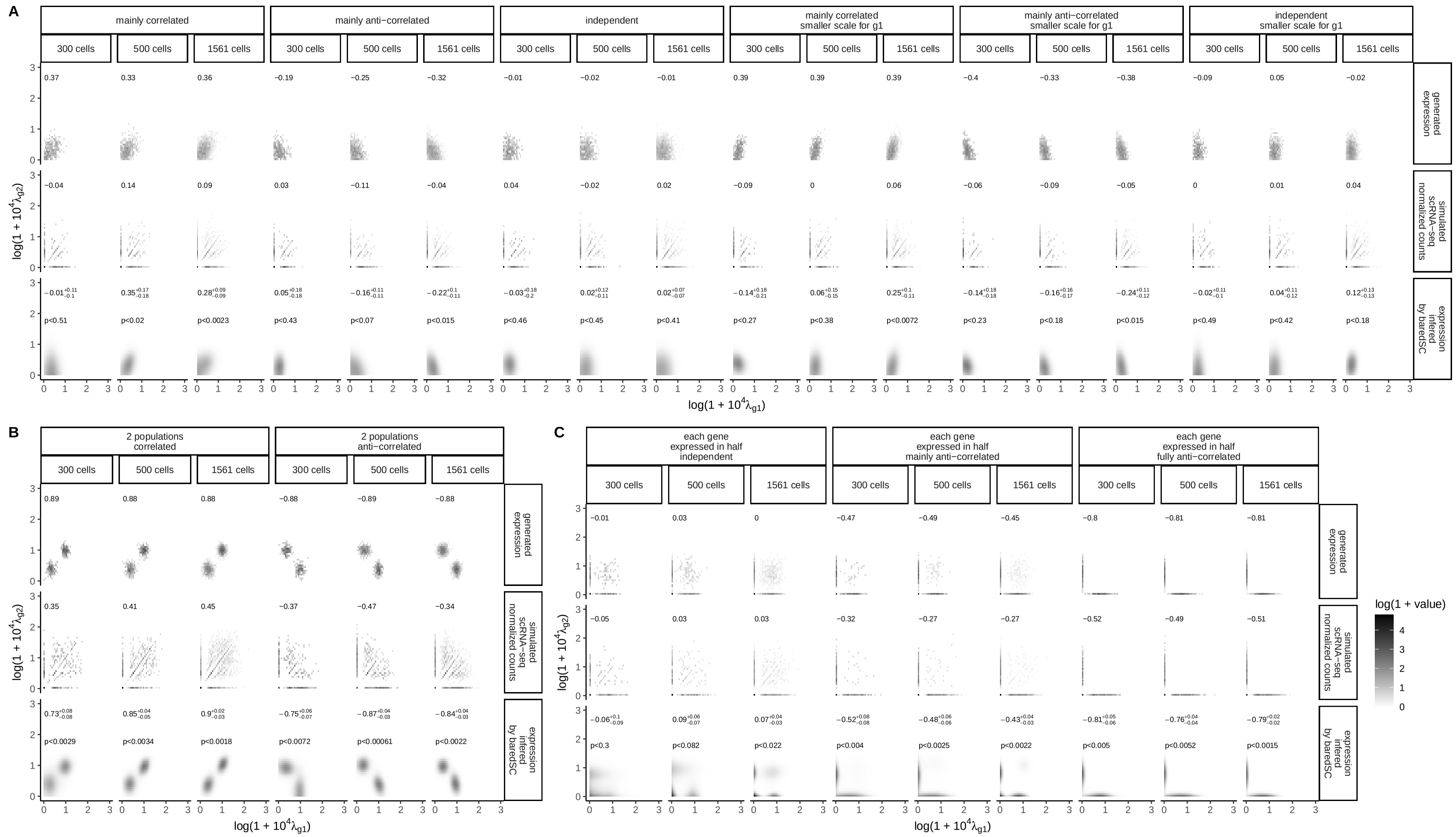


**Supplementary Fig. 2.2.2: baredSC in 2D with smaller number of cells.** This figure is identical to Figure 3 except that the result for each group of cells is plotted.

#### 2.3 Analysis of distribution of *Pitx1* expression in log axis

In baredSC, we implemented two scales. The scale used in the Seurat package which is $\log(1 + 10^4 X_{i,g})$ and the regular log scale ($\log(X_{i,g})$). In all main figures of this article except Fig. 3, we used the Seurat scale. This choice is motivated by the decreased degeneracy of models in Seurat scale for cells with no count compared to the log scale. However, the log scale may be more convenient in some cases.

In particular, we use it in Fig. 3 and Supplementary Figs. 2.3.3 to compare to other techniques which use the log scale. In the case of *Pitx1* expression, we have fluorescence levels from flow cytometry which are conventionally represented in log scale (Supplementary Fig. 2.3.3A). We use baredSC to retrieve the PDF in log scale (Supplementary Fig. 2.3.3B). For FL *Pitx1*$^{+/+}$ (in black), where *Pitx1* is not expressed, we observe that the distribution is concentrated on the low values. However, we observe a larger confidence interval compared to the distribution obtained in the Seurat scale (Fig. 5D). This is due to the degeneracy of the model at low value of expression: having no count is similarly probable for expression values of -13 in log scale (2e-6 which is much smaller than $1/N_i$) or below. Thus, one should consider the low expression range as cells that do not express *Pitx1*. As explained in the article, in this range of expression, it is difficult to compare both techniques as the flow cytometry has a high level of background. At higher level of expression (right part), the PDF of HL *Pitx1*$^{Pen-/Pen-}$ and HL *Pitx1*$^{+/+}$ have a highly similar shape between the flow cytometry (Supplementary Fig. 2.3.3A) and the baredSC results (Supplementary Fig. 2.3.3B). These results confirm what we observed in Fig. 5 that the fluorescence level is highly correlated to the level of expression of *Pitx1*. We then compared these results to the density plot of normalized expression (Supplementary Fig. 2.3.3C). For this representation of the normalized counts, the expression of cells with no expression was artificially put to the minimal value. We see that the PDF of cells where *Pitx1* is detected is close to a Gaussian for both HL samples. We also run Sanity to visualize the posterior distribution given by this algorithm (Supplementary Fig. 2.3.3D). We see that the tissue which does not express *Pitx1* (FL *Pitx1*$^{+/+}$ in black) is described as a large Gaussian, suggesting that a fraction of cells expresses *Pitx1* at low level. The PDF of samples which express *Pitx1* (HL in blue and red) is unimodal. This show that the posterior distribution from Sanity is not able to detect the trimodal expression of *Pitx1* in HL samples.

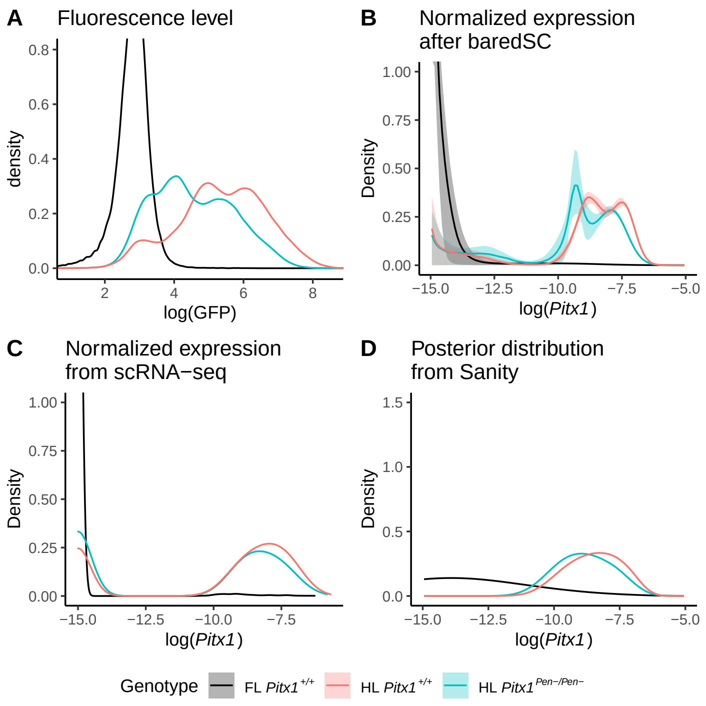


**Supplementary Fig. 2.3.3: baredSC retrieves the trimodal distribution of *Pitx1*.** **A**: Distribution of fluorescence intensity obtained by flow cytometry in regular log scale. **B**: Distribution of normalized expression from scRNA-seq as provided by baredSC when run in log scale. **C**: Distribution of normalized expression from scRNA-seq with a regular log scale. **D**: Posterior distribution obtained with Sanity.

### 3 Bibliography

#### References

Andrieu, C. and Thoms, J. (2008). A tutorial on adaptive mcmc. *Statistics and Computing*, 18(4):343–373.

Delisle, J.-B., Ségransan, D., Dumusque, X., Diaz, R. F., Bouchy, F., Lovis, C., Pepe, F., Udry, S., Alonso, R., Benz, W., Coffinet, A., Cameron, A. C., Deleuil, M., Figueira, P., Gillon, M., Curto, G. L., Mayor, M., Mordasini, C., Motalebi, F., Moutou, C., Pollacco, D., Pompei, E., Queloz, D., Santos, N. C., and Wyttenbach, A. (2018). The HARPS search for southern extra-solar planets. XLIII. A compact system of four super-Earth planets orbiting HD 215152. *A&A*, 614:A133.

Haario, H., Saksman, E., and Tamminen, J. (2001). An adaptive metropolis algorithm. *Bernoulli*, 7(2):223–242.
